# Drivers of immune-related genetic variation across human populations

**DOI:** 10.64898/2026.06.29.735156

**Authors:** Adriana Morales-Guerrero, Keith D. Harris, Nimrod Marom, Gili Greenbaum

## Abstract

Pathogen-mediated disease burden imposes some of the strongest selective pressures on human populations, shaping genetic variation in immune-related genomic regions. Because historical, cultural, ecological, and environmental factors can influence disease burden, associations between these factors and immune-related genomic signatures can provide insight into how pathogen-mediated selection varied among populations. To investigate these associations and identify factors potentially contributing to historical disease burden, we analyzed worldwide variation in the Major Histocompatibility Complex (MHC). We evaluated the relationships between global patterns of genetic variation and a comprehensive set of historical-cultural, ecological, and climatic variables compiled in DisECCO (Disease, Ecological, Cultural, and Climatic Origins dataset). This dataset includes factors potentially associated with historical disease burden, including the timing of major cultural transitions, historical climatic conditions, exposure to domesticated animals, and pre-industrial pathogen stress. Contrary to genome-wide expectations, MHC diversity and variation were not well explained by geographic distance from Africa, suggesting that neutral demographic processes play a limited role in shaping MHC variation. Instead, MHC genetic variation was significantly associated with climatic variables, domestication exposure, and the time lag between the onset of the Neolithic and urbanization, with the strongest explanatory models identified for MHC Class II. These results indicate that cultural transitions and environmental conditions have played a central role in shaping immune-related genetic variation, and may have contributed substantially to changes in disease burden. Overall, our findings show that integrating ecological and cultural-historical variables with genomic data can help explain global patterns of genetic variation in humans.

## Introduction

Pathogens have imposed a substantial burden on human populations worldwide [1], exerting strong selection pressures [2, 3]. The intensity of pathogen-mediated selection depends on the historical levels of pathogen exposure and transmission, which have been shaped by a combination of cultural, ecological, demographic, and climatic factors. Consequently, pathogen-driven selection pressures have varied across populations and through time, leading to differences in immune-related genomic signatures among modern populations [4]. Historical disease burden is therefore not only a driver of immune evolution, but may also reflect historical transitions in human ecology and culture [5, 6]. However, the extent to which variation in immune-related genomic signatures reflects historical variation in disease burden, and the factors that have shaped these patterns, remain unclear.

Pathogen-induced disease burden can leave genetic signatures through different selective mechanisms. While a single disease or epidemic typically induces directional selection that favors alleles conferring resistance (e.g., [7–11]), the cumulative burden of multiple diseases over long timescales can promote balancing selection and maintain genetic diversity in immune-related genomic regions [12, 13]. This balancing selection is mainly attributed to two mechanisms: heterozygote advantage in immune-related regions, where broader immune repertoires enhance pathogen recognition, and negative frequency-dependent selection, where rare alleles are advantageous because pathogens are less likely to have evolved evasion strategies against them [12]. As a result of this balancing selection and the strong selective pressures imposed by pathogenic environments, immune-related genomic regions are among the most genetically diverse in the human genome [14].

Genetic signatures of disease burden are most evident in the Major Histocompatibility Complex (MHC), the central component of the immune system, which presents protein fragments to lymphocytes allowing the detection of pathogens and the activation of immune responses [12, 15]. The MHC comprises three major sub-regions, each associated with distinct immune functions: MHC Class I, which presents endogenous antigens and is primarily involved in responses to viruses [16]; MHC Class II, which presents exogenous antigens and is primarily linked to responses to bacterial infections [17, 18]; and MHC Class III, which encodes a diverse range of proteins involved in innate and adaptive immune responses, particularly inflammation and post-transcriptional gene regulation [19]. Variation in MHC-I and MHC-II, which directly initiate adaptive immune responses, is expected to reflect pathogen-specific selection more strongly than MHC-III, which has genes that are involved in broader and non-specific immune functions. Because these classes participate in distinct immune pathways and respond to different categories of pathogens, they may differ in the extent to which their genetic variation reflects historical pathogen-mediated selection.

Genome-wide variation among human populations is fairly well understood, with diversity primarily shaped by geographic distance from Africa along human migration routes [20, 21]. Immune-related regions such as the MHC, however, are expected to depart from this neutral demographic background because they are also shaped by pathogen-mediated selection. Consistent with this expectation, population-level variation in the MHC has been linked to disease ecology variables, particularly differences in pathogen richness and pathogen load [2, 22, 23]; for example, a comparative study of African hunter-gatherers and agriculturalists showed differential immune responses driven mainly by viral exposures [24]. These studies support the idea that MHC variation can reflect variation in disease burden, while also highlighting a key limitation: most available predictors capture present-day or recent differences in living conditions, rather than the historical cultural, ecological, and environmental processes that shaped pathogen exposure over evolutionary timescales.

Major cultural transitions are among the clearest examples of how human history may have reshaped disease burden by modifying pathogen exposure [5, 6]. The Neolithic shift to a sedentary agricultural lifestyle, for example, led to higher population densities and increased connectivity, potentially facilitating the spread and persistence of infectious diseases [5, 25, 26]—an idea recently supported by analyses of ancient human [27] and pathogens genomes [28]. The later emergence of urbanism likely augmented this effect [29, 30], as did the widespread adoption of domesticated animals, which intensified human exposure to zoonotic pathogens and thereby increased the disease burden experienced by populations [25, 31]. In parallel, spatial and temporal variation in climatic conditions may also have shaped pathogen landscapes, influencing patterns of disease burden across regions [32, 33]. Therefore, variability in the timing, intensity, and characteristics of cultural and climatic factors likely contributed to shaping immune-related selection pressures across populations. However, how this variability contributed to disease burden and thereby shaped selective pressures on MHC variation, and what was the relative importance of the different factors, remains unclear.

Here, we evaluate selective drivers of genetic variation in the MHC region across contemporary human populations using a newly compiled Disease, Ecological, Cultural, and Climatic Origins (DisECCO) dataset, which integrates a wide spatio-temporal range of variables. This dataset enabled us to assess how diverse factors are associated with immune-related signatures of balancing selection in the MHC as a whole and within its three major sub-regions (Classes I, II, and III). Through this analysis, we identify specific cultural and environmental predictors that best explain immune-related genomic variation in humans and highlight their potential role in shaping pathogen-mediated selection in the MHC.

## Results

### Patterns of genetic variation across MHC regions

To study genetic diversity and balancing selection in the MHC, we used the Human Genome Diversity Project (HGDP) dataset, a geographically distributed genomic resource comprising 929 individuals from 54 relatively unadmixed populations [34, 35]. Because the HGDP captures human populations spanning a wide range of cultural, ecological and climatic contexts while preserving signals of deeper demographic history, it provides an opportunity to examine how historical drivers of disease burden are associated with contemporary patterns of immune-related genetic variation. The dataset includes high-resolution genomic data of > 67 million single nucleotide polymorphisms (SNPs), about 158,000 of which are located within the MHC region. We evaluated two measures of genetic variation that reflect different aspects of balancing selection: (i) genetic diversity, measured as the mean observed heterozygosity of each population, where stronger balancing selection on a genomic region is expected to result in higher heterozygosity levels [36]; (ii) Tajima’s *D* [37], where positive values indicate an excess of intermediate-frequency alleles that is associated with balancing selection, but could also arise as a result of demographic processes. We measured these two statistics for the MHC region as a whole (Fig. 1A and Fig. 1B), in addition to a partition of the MHC into the three main functional regions (denoted as MHC-I, MHC-II, and MHC-III; Fig. S1). Among these sub-regions, MHC-II exhibited significantly higher Tajima’s *D* and heterozygosity than both MHC-I and MHC-III (paired Wilcoxon tests; Fig. S2). As control, we also calculated both measures across the genome, excluding the MHC region (Fig. S3). For mean observed heterozygosity, we quantified excess genetic diversity in the MHC and its sub-regions by subtracting genome-wide heterozygosity from the heterozygosity measured in each region (Δ*H_o_*). Unless otherwise stated, all references to heterozygosity in the MHC refer to this measure.

**Figure 1:**
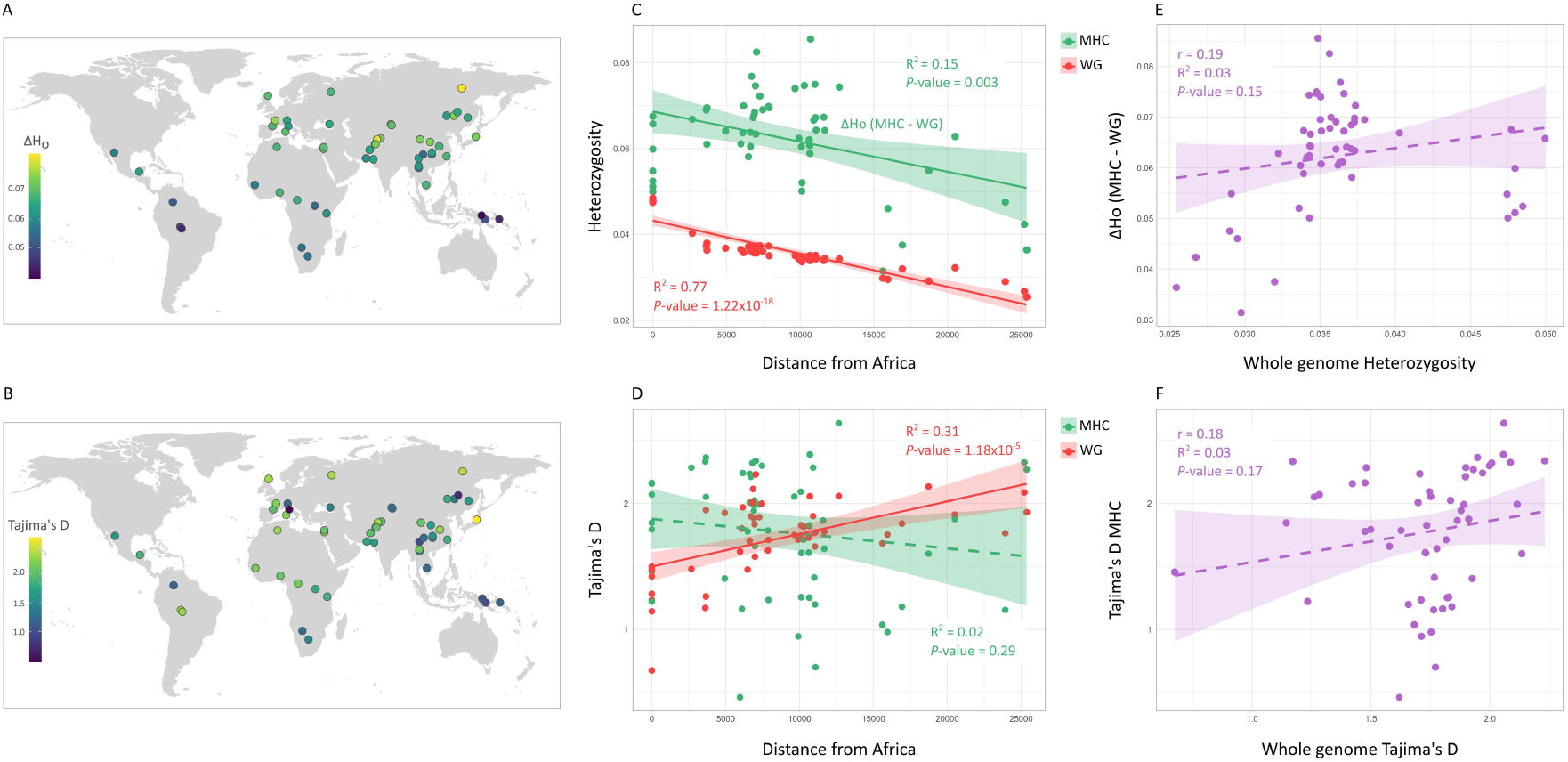
Worldwide patterns of genomic variation in the MHC relative to the whole genome. All analyses were conducted on 54 populations from the HGDP. (A) Excess MHC heterozygosity (Δ*H_o_*) for each population. (B) MHC Tajima’s *D* of each population. (C–D) Associations of heterozygosity and Tajima’s *D* with distance from Africa, where distances are constrained through waypoints to reflect human migration routes. MHC measures are shown in green, and genome-wide measures in red. Colored lines and text show results of linear regressions, with continuous lines denoting significant associations and dashed lines non-significant ones. Shaded areas show the 95% confidence intervals. (E–F) Relationships between MHC and genome-wide values of heterozygosity (E) and Tajima’s *D* (F). Purple dashed line denotes non-significant linear regressions.

### MHC variation is not well-explained by distance from Africa

One of the strongest predictors of genome-wide variation among human populations is geographic distance from Africa, measured along human migration routes [20, 21]. In our analysis, this pattern is reflected in a strong and significant correlation between genome-wide genetic diversity and distance from Africa (*R*^2^ = 0.77, *r* = −0.88, *P* = 1.22 × 10^−18^; red in Fig. 1C). genome-wide Tajima’s *D* signal was also significantly and positively correlated with distance from Africa (*R*^2^ = 0.31, *r* = 0.56, *P* = 1.19 × 10^−5^; red in Fig. 1D), but to a lesser extent than heterozygosity. This positive association reflects the excess of intermediate-frequency variants in populations farther from Africa, caused by repeated founder effects along human migration routes and the consequent loss of rare alleles. We consider these patterns as the “neutral” demographic background against which the measures of genetic variation in the MHC can be compared.

We then evaluated the relationship between MHC genetic variation across populations and their distance from Africa. In contrast to the genome-wide analysis, we found no association between MHC Tajima’s *D* and the distance from Africa (green in Fig. 1D), and only a weak association for MHC heterozygosity (*R*^2^ = 0.15, *r* = −0.38, *P* = 0.03; green in Fig. 1C). We also found that the measures of genetic variation in MHC-I and MHC-III were not significantly associated with distance from Africa, while those of MHC-II were weakly associated (Table S1).

To further examine the extent to which MHC patterns are shaped by processes different from those shaping genome-wide variation, we evaluated the relationship between genome-wide and MHC diversity. No significant associations were found when measuring heterozygosity (Fig. 1E) or Tajima’s *D* (Fig. 1F). Moreover, we found no significant correlation between the genome-wide measures of genetic variation and those of any of the three MHC functional sub-regions (Tables S2 and S3). By contrast, we did find many associations among the MHC classes themselves (Tables S2 and S3), indicating that variations in MHC sub-regions are more similar to one another than to the genome as a whole. Although the slopes of the linear regressions in Figure 1E–F were not significant, the intercepts for both heterozygosity and Tajima’s D relationships were significantly positive (intercept of MHC heterozygosity vs. genome-wide heterozygosity = 0.05, *P* = 4.17 × 10^−5^; Intercept of MHC Tajima’s D vs. genome-wide Tajima’s D = 1.21, *P* = 5.5 × 10^−3^), indicating a consistently elevated heterozygosity and an excess of intermediate-frequency alleles in the MHC relative to genome-wide expectations, patterns that are consistent with immune-related balancing selection. Altogether, these results suggest that, although genome-wide neutral variation is strongly shaped by the migration out-of-Africa, substantial variation in the MHC remains unexplained and may be better accounted for by considering additional historical–cultural, ecological, and climatic predictors beyond distance from Africa.

### Curating historical predictors of disease burden for studying immune-related genomic variation

Although genomic datasets such as the HGDP provide a powerful resource for studying variation among human populations, they do not include the historical-cultural, ecological, and climatic information needed to evaluate hypotheses about historical disease burden. Many contextual factors could plausibly be associated with disease-burden related genomic signatures, including cultural transitions [24, 26], pathogen exposure [2, 22], domestication history, climate [38], and demographic history [39]. However, these variables must first be collected, standardized, and assigned to populations in a consistent framework. We, therefore, curated a set of predictors designed to capture historical processes that may have shaped pathogen exposure, transmission, intensity, and disease burden across human populations. Because MHC variation was not well explained by either distance from Africa or genome-wide variation, this framework allowed us to test whether historical disease-burden predictors could explain immune-related genomic variation beyond broad neutral demographic patterns.

The DisECCO dataset comprises a comprehensive set of predictor variables that pair HGDP populations with historical-cultural, ecological, and climatic factors: (i) pre-industrial pathogen stress estimates, (ii) the timing of two major cultural transitions, the onset of Neolithic lifestyle and the onset of urbanization, (iii) the time lag between these two transitions (Δ*N* –*U*), (iv) climatic conditions, both current and during each major transition, (v) a sociopolitical complexity index integrated over time, and (vi) overall exposure to domesticated animals (Fig. 2). Together, these variables provide a contextual framework for studying how historical processes may be associated with genomic variation across human populations. In this study, we use this framework to evaluate whether proxies for historical disease burden and related cultural-ecological processes can explain immune-related genomic variation. Several predictors in DisECCO, including pathogen stress, climatic conditions, and exposure to domesticated animals, have previously been linked to pathogen exposure or disease burden and were therefore considered plausible candidates for explaining immune-related genomic variation. Others, including sociopolitical complexity and the lag between the onset of the Neolithic and urbanization (Δ*N* –*U*), provide an opportunity to evaluate less explored hypotheses regarding the historical drivers of immune-related genomic variation.

**Figure 2:**
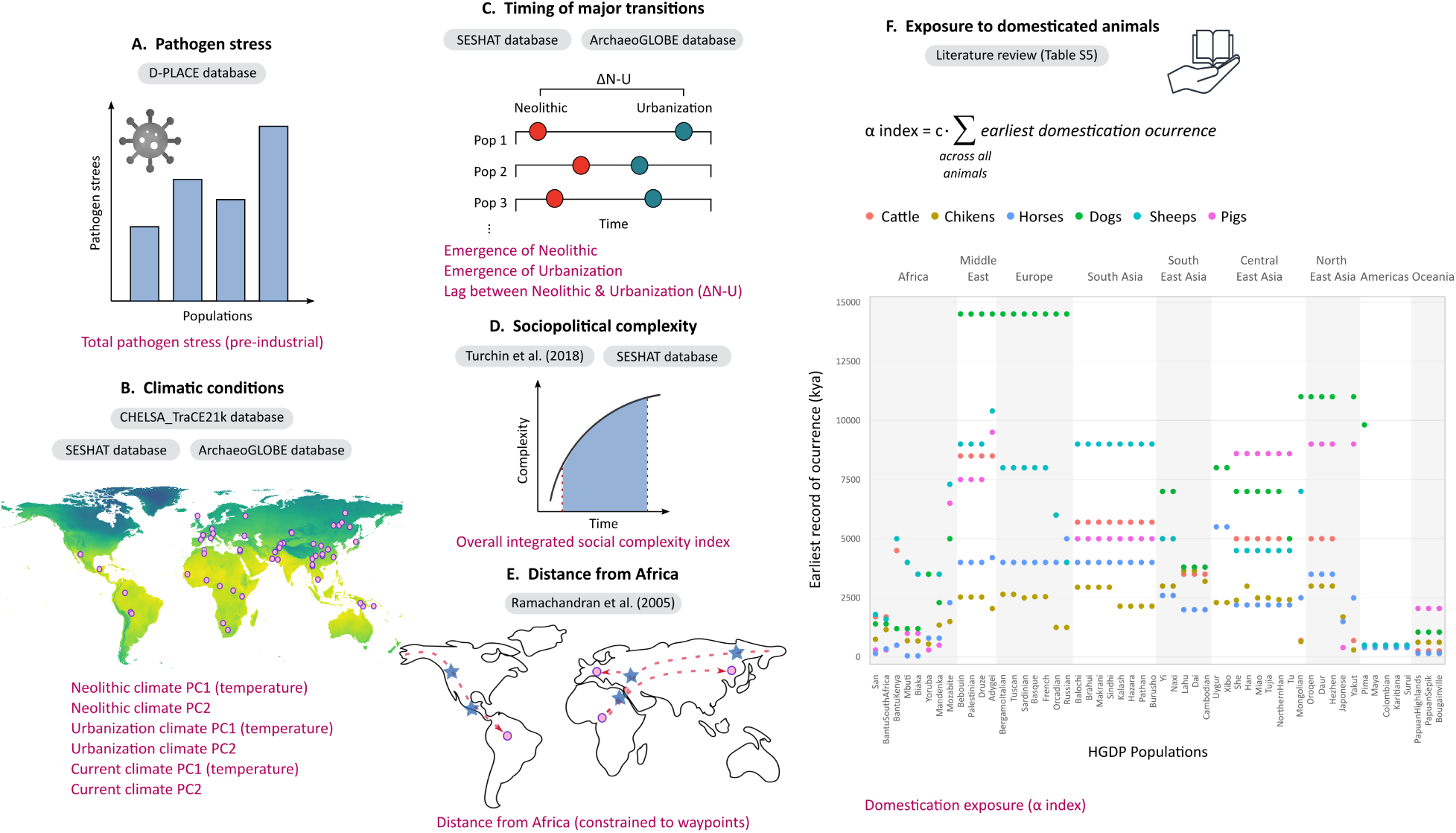
Overview of the DisECCO dataset. All predictor variables were assigned to each of the HGDP populations. (A) Pre-industrial pathogen stress, defined as the total pathogen stress from the D-PLACE database [44]. (B) Climatic conditions at three time points: current day, onset of Neolithic, and onset of urbanization. The timing of the two major historical transitions was determined using the Seshat and ArchaeoGLOBE datasets [40, 41]. Climatic variables for each time point were extracted from the CHELSA-TraCE21k database [43], and summarized by the first two principal components of 19 bioclimatic variables within each region. (C) Timing of two major transitions, the Neolithic and urbanization, as defined in the Seshat and ArchaeoGLOBE datasets [40, 41]. We defined Δ*N* –*U* to denote the lag between these two major transitions. (D) Sociopolitical complexity index, defined as the area under the curve (AUC) of the first principal component from Turchin *et al.* [42], over time. (E) Distance from Africa, constrained to pass through known human migration waypoints. (F) Exposure to domesticated animals, defined using an index, *α*, which quantifies the exposure to domesticated animals over time, based on the earliest domestication events per domesticated species. The data for calculating *α* was collected from available literature (shown here is a subset of the data; the full data is available in *Supplementary Dataset S4*).

We first compiled predictors describing major cultural transitions and long-term sociopolitical change. We assessed the timing of the onset of Neolithic lifestyle and urbanization by integrating information from large-scale historical datasets and literature sources [40, 41], assigning estimates to each HGDP population according to the closest geographic region (*Supplementary Dataset S1*). Based on these estimates, we calculated the time lag between the two transitions (Δ*N* –*U*) for each population (Fig. 2C). We also assessed sociopolitical complexity using a dataset summarizing 51 sociopolitical variables over the last 10,000 years for 30 geographic regions [42]. We used PC1 of these variables and calculated the area under the curve (AUC) of PC1 over time for each region as an estimate of overall sociopolitical complexity, which was then assigned to HGDP populations based on geographic proximity (*Supplementary Dataset S2*).

We also compiled predictors describing climatic conditions and pathogen-exposure related processes. Climatic conditions were summarized using a principal component analysis (PCA) integrating current climatic variables from the CHELSA v2.1 bioclimatic dataset with historical multivariate climatic data from the CHELSA-traCE21k database [43]. For historical estimates, climatic conditions were extracted for the geographic location associated with each population at the onset of the Neolithic and the emergence of urbanization. We used the first two principal components, PC1 and PC2, which explained 52% and 18% of the climatic variance, respectively (*Supplementary Dataset S3*); PC1 was primarily related to temperature variation, while PC2 mainly captured variation in precipitation (Fig. S4). To quantify exposure to domesticated animals, we conducted an extensive literature review (Fig. 2F, *Supplementary Dataset S4*), compiling estimates of the timing of introduction of numerous domesticated species across the geographic regions associated with the 54 HGDP populations. We then developed an index, *α*, which accounts for both the duration of exposure of each population to domesticated animals and the number of domesticated animals. Finally, to assess pre-industrial pathogen stress, we matched each HGDP population to the geographically closest non-industrial population represented in the D-PLACE database [44] (*Supplementary Dataset S5*).

Full definitions and assignment procedures are provided in *Methods* and the complete DisECCO dataset is available in *Supplementary Dataset S6*. Correlations between the predictors are shown in Figure S5; the strongest correlations were observed among the climatic predictors (|*r*| ≥ 0.8).

### Key cultural, ecological, and climatic predictors explain MHC variation

Using the DisECCO dataset, we performed a multivariate linear regression analysis to evaluate the relationship between each measure of genetic variation and the set of predictors. We also included geographic distance from Africa as a control predictor to account for neutral demographic patterns associated with human migration history. Analyses were performed separately for the whole MHC, for each of the three MHC classes, and for the genome-wide as a control. We identified the best–fitting models using LASSO and AICc criteria and assessed collinearity among best predictors by calculating the mean variance inflation factor (*V IF*) [45], with values close to one indicating low collinearity. To assess the robustness of our findings, we conducted a sensitivity analysis by progressively introducing random errors into the 13 predictors and re-running the regression analysis. The contribution of a predictor to explaining genomic patterns was considered robust if it was retained in at least 50% of the best–fitting models under up to 20% random error (Figs. S6–S7).

In the MHC, the best–fitting model for Tajima’s *D* retained three predictors that together explain 18.7% of the variance (*R*^2^ = 0.187, *P* = 0.015; Fig. 3A, Fig. S6A). This model had a low multicollinearity factor (*V IF* = 1.03), indicating that the predictors in the model are not substantially correlated with each other. The onset of urbanization predictor accounted for the largest proportion of explained variance (7%), followed by the lag between the major transitions Δ*N* –*U* (6%), and pre-industrial pathogen stress (5%). The Δ*N* –*U* predictor was negatively associated with Tajima’s D, indicating that shorter lag times were associated with higher Tajima’s *D* values, (i.e., stronger balancing selection signal for shorter lag times). The other two predictors were positively associated with Tajima’s *D*, meaning that higher pre-industrial pathogen stress and earlier onset of urbanization were associated with stronger signatures of balancing selection in the MHC. For MHC genetic diversity, three predictors were retained in the best–fitting model (*R*^2^ = 0.610, *P* = 2.5 × 10^−10^, *V IF* = 1.42; Fig. 3B, Fig. S6B). Here, PC1 of Neolithic climatic conditions explained the largest proportion of variance (38%), followed by pre-industrial pathogen stress (13%) and the *α* index (10%). The association between PC1 of Neolithic climatic conditions and heterozygosity was negative, indicating that colder temperatures are associated with higher levels of MHC diversity. Pre-industrial pathogen stress was positively associated with MHC diversity, as was our *α* index for the exposure to domesticated animals.

**Figure 3:**
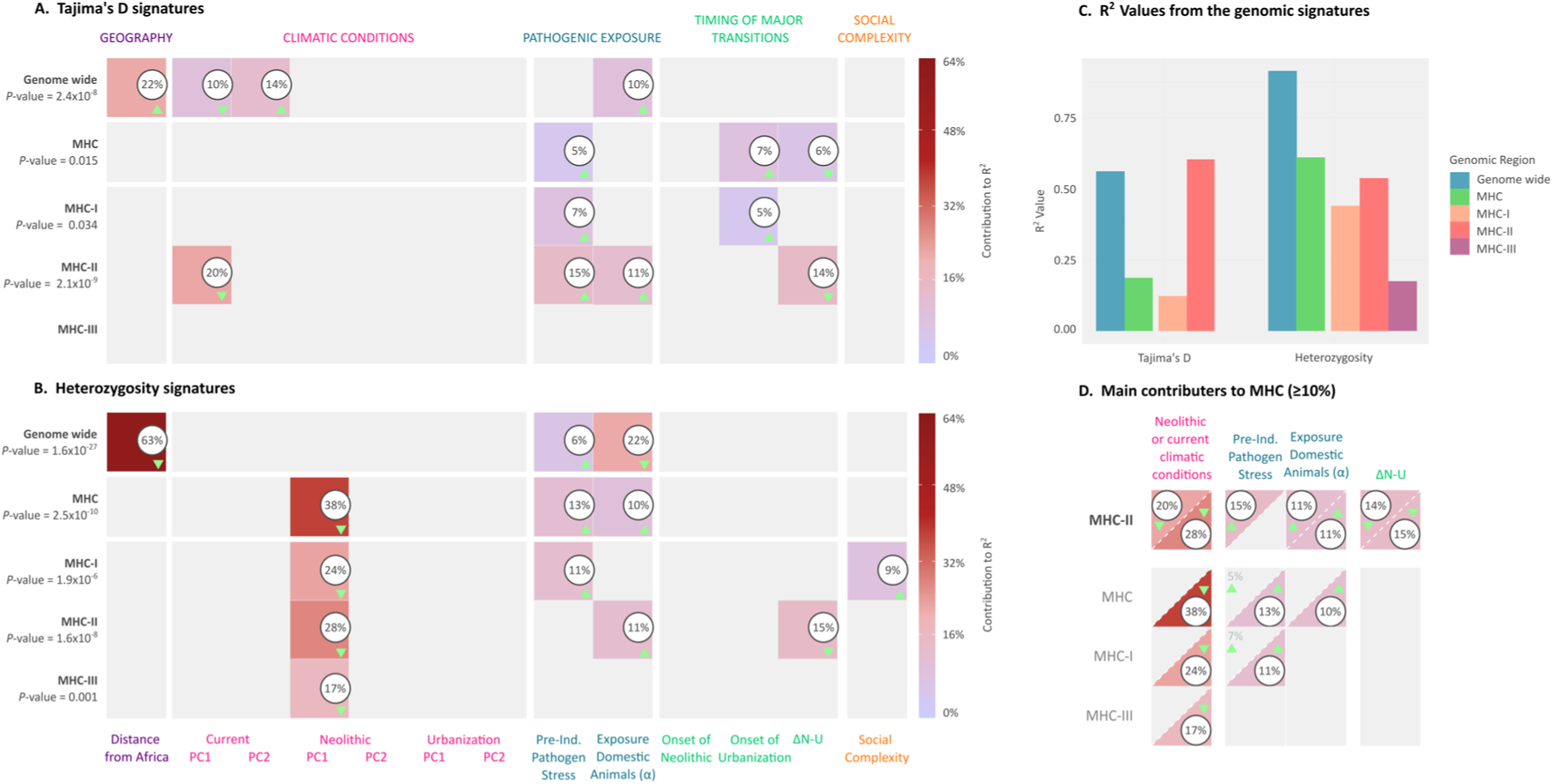
Best–fitting multiple regression models explaining worldwide immune-related genetic variation. (A) Best–fitting models explaining variation in Tajima’s *D* values across 54 populations. (B) Best–fitting models explaining variation in mean heterozygosity values across 54 populations. In A and B, each row corresponds to single model focusing on a different genomic region, with *p* − *values* indicated on the left. Colored squares indicate the relative contribution of each predictor to the model’s explained variance, with darker shades representing higher contributions. Numbers within squares denote the portion of the variance explained by each predictor that is included in the model. Small green arrows within the colored squares indicate the direction of the relationship: upward for positive, downward for negative. (C) Comparison of the explanatory power (*R*^2^) of the best–fitting models for each genomic region and genetic measure. Bars represent the proportion of variance explained by the corresponding multiple regression model. (D) Predictors retained in the best-fitting MHC-II models, each explaining more than 10% of the variance. The corresponding contributions of these predictors in other MHC classes are shown below when they also exceeded this threshold. Within each square, values above the diagonal correspond to Tajima’s *D*, whereas values below the diagonal correspond to heterozygosity.

When focusing on genetic variation in MHC-I, the best–fitting model for Tajima’s D included only two predictors (*R*^2^ = 0.123, *P* = 0.034, *V IF* = 1.00; Fig. 3A, Fig. S6C), pre-industrial pathogen stress (explaining 7% of the variation) and the onset of urbanization (explaining 5% of the variation). Both predictors were also included in the model for the whole MHC, with the same direction of the associations. For the genetic diversity in MHC-I, the best–fitting model retained three predictors (*R*^2^ = 0.440, *P* = 1.9 × 10^−6^, *V IF* = 1.60; Fig. 3B, Fig. S6D), PC1 of Neolithic climatic conditions (24% of the variation), pre-industrial pathogen stress (11% of the variation) and the sociopolitical complexity (9%). As in the overall MHC model, MHC-I diversity was negatively associated with PC1 of Neolithic climatic conditions, which primarily reflects temperature, and positively associated with pre-industrial pathogen stress. Sociopolitical complexity was also positively related to MHC-I diversity, suggesting that greater complexity corresponds to higher levels of MHC-I diversity.

The best–fitting models for MHC-II were substantially better than for the other MHC classes (Fig. 3C), particularly for Tajima’s *D*, where we were able to explain more than 60% of the variation using the DisECCO dataset (compared to 18% for the whole MHC or 56% for the genome-wide variation). The best–fitting model for MHC-II Tajima’s *D* retained four predictors (*R*^2^ = 0.604, *P* = 2.1 × 10^−9^, *V IF* = 1.52, Fig. 3A, Fig. S6E): PC1 of the current climatic conditions (explaining 20% of the variation), pre-industrial pathogen stress (15%), Δ*N* –*U* (14% of the variation), and the exposure to domesticated animal index *α* (11% of the variation). Both PC1 of current climatic conditions and Δ*N* –*U* showed negative associations, suggesting that colder environments and shorter intervals between the onset of the Neolithic and urbanization are linked to stronger balancing selection in MHC-II, whereas the remaining predictors showed positive effects. The best model for MHC-II genetic diversity was also better than the models for the diversity in other sub-regions, explaining more than 50% of the variation in heterozygosity, similar to the model for diversity in the whole MHC (Fig. 3C). The best–fitting model for MHC-II diversity included three predictors, (*R*^2^ = 0.538, *P* = 1.6 × 10^−8^, *V IF* = 1.44; Figure 3B, Fig. S6F): PC1 of the Neolithic climatic conditions accounted for the largest proportion of explained variance (28% of the variation), followed by the Δ*N* –*U* (15% of the variation), and the *α* index (11% of the variation). As with Tajima’s D, both PC1 of the Neolithic climatic conditions and Δ*N* –*U* were negatively associated with heterozygosity, indicating that colder environments and shorter transition periods were linked to increased heterozygosity. The *α* index was positively associated with MHC-II diversity and with Tajima’s *D*, indicating an association between higher levels of exposure to domesticated animals and stronger signatures of balancing selection in MHC-II.

In MHC-III, on the contrary, we could not identify any model or variable that met the robustness criterion to explain the variation in Tajima’s D (Figure 3C, Fig. S7A); we only identified a weak negative association between heterozygosity and PC1 of the Neolithic climatic conditions (*R*^2^ = 0.175, *P* = 0.001, Fig. S7B). This is perhaps expected given the more indirect and mediated role of MHC-III in pathogen-related immune responses.

We also examined the relationship between the set of predictors and the genome-wide Tajima’s *D* and heterozygosity, as a control for our immune-related models. In contrast to the immune-related regions, genome-wide variation was primarily shaped by geographic distance from Africa along the out-of-Africa migration routes, reflecting the associations shown in Figure 1C and D. For genome-wide Tajima’s *D*, the best–fitting model retained four predictors (*R*^2^ = 0.562, *P* = 2.4 × 10^−8^, *V IF* = 1.34, Fig. 3A, Fig. S7C). Distance from Africa had the strongest effect, explaining 22% of the variance, with PCs 1 and 2 of current climate (10% and 14%, respectively) and the *α* index (10%) contributing as well. The best–fitting model for genome-wide diversity explained more than 90% of the variation (*R*^2^ = 0.915, *P* = < 1.6 × 10^−27^, *V IF* = 1.18; Figure 3B, Fig. S7D). This model included three predictors: distance from Africa, which explained 63% of the variance, as well as the *α* index (22%; notably, the association here is negative, the opposite from that observed in the MHC classes), and the pre-industrial pathogen stress (6%). The high explanatory power of these two models (Figure 3C), primarily driven by distance from Africa, highlights the dominant influence of migration in shaping genome-wide diversity, in contrast to the more nuanced and culturally mediated patterns observed in immune-related genomic regions.

In summary, among the three MHC classes, MHC-II showed the highest proportion of variance explained by our DisECCO dataset (Fig. 3C) and also exhibited stronger signals of balancing selection and higher genetic diversity across populations than MHC-I and MHC-III (Fig. S2). Considering both measures of genetic variation, four key predictors emerged as the contributors to MHC-II variation, each accounting for more than 10% of the explained variance (Fig. 3D): (i) climatic conditions during either the current (for Tajima’s *D*) or Neolithic (for heterozygosity) periods, (ii) exposure to pathogens, captured by pre-industrial pathogen stress and long-term contact with domestic animals (*α* index), and (iii) the temporal lag between the onset of agriculture and urbanization (Δ*N* –*U*). Several of these predictors also exceeded this threshold in other MHC classes (Fig. 3D), suggesting that they represent key factors shaping immune-related genomic variation.

### Climate, time-lag between major transitions, and exposure to pathogens drive variation in MHC class II

To further evaluate independent effects of the main predictors, we conducted univariate regressions between each of the four key predictors of MHC-II variation and the corresponding measures of genetic variation. Consistent and significant associations were observed for three of these predictors (climatic conditions, the temporal lag between the major cultural transitions, and long-term domestication exposure), indicating that their relationships with MHC-II variation were detectable even when considered individually (Fig. 4). For current and Neolithic climatic conditions, the associations were very strong (Fig. 4A–B) and explained a considerable portion of the overall variance: Current climate accounted for 36% of the variation in Tajima’s *D* (*R*^2^ = 0.359, *P* = 1.6 × 10^−6^; Fig. 4A), while Neolithic climate explained 47% of the variation in heterozygosity (*R*^2^ = 0.472, *P* = 9.4 × 10^−9^; Fig. 4B). In both cases, and consistent with the multivariate models, colder and temperate climates were associated with higher Tajima’s *D* and heterozygosity in MHC-II, patterns consistent with stronger balancing selection. Populations exhibiting these signatures were primarily located in Central and North-East Asia, whereas populations with lower heterozygosity and Tajima’s *D* values were mainly found in equatorial regions of the Americas, Africa, and Oceania (Fig. 4C).

**Figure 4:**
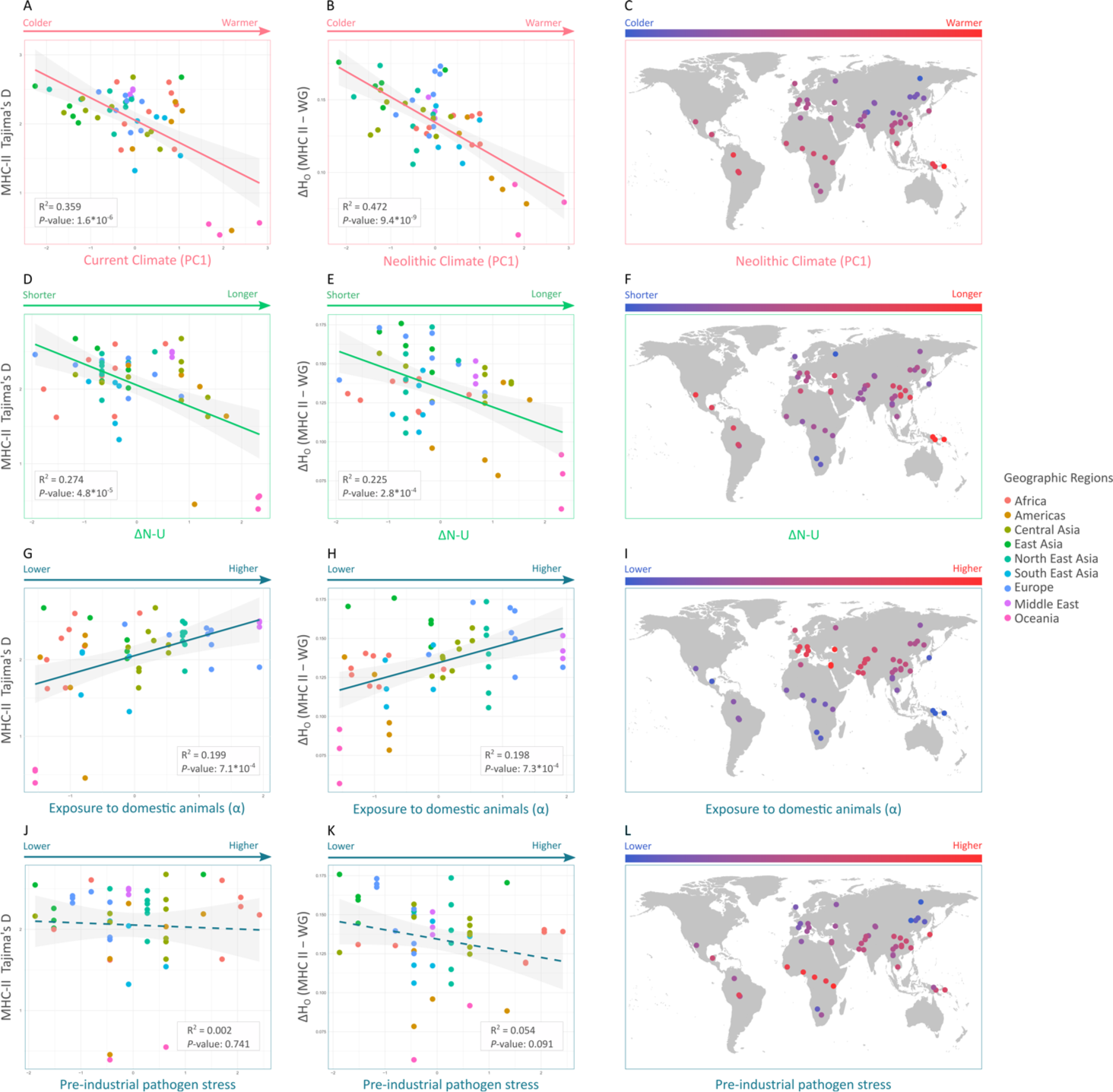
Association between the MHC-II contributing predictors and the measures of genetic variation. Shown are the four contributor predictors in the multivariate best-fitting models, for both Tajima’s *D* and heterozygosity in the MHC-II sub-region. Each predictor is shown in an independent regression to evaluate its direct association with each genomic signature. (A) Relationship between current climate (PC1) and MHC-II Tajima’s *D*. (B) Relationship between Neolithic climate (PC1) and MHC-II heterozygosity. (C) Geographic distribution of the Neolithic climate (PC1) predictor across the 54 HGDP populations. Circle colors indicate climate values, with warmer colors representing higher temperatures and cooler colors representing lower temperatures. (D–E) Relationships between Δ*N* –*U* and MHC-II Tajima’s *D* and heterozygosity. (F) Geographic distribution of Δ*N* –*U*. Circle colors indicate transition lengths, with warmer colors representing longer lags between the Neolithic and urbanization and cooler colors shorter lags. (G–H) Relationships between domestication exposure (*α*) and MHC-II Tajima’s *D* and heterozygosity. (I) Geographic distribution of *α*. Circle colors indicate exposure levels, with warmer colors representing higher exposure to domestic animals over time and cooler colors lower exposure. (J–K) Relationships between pre-industrial pathogen stress and MHC-II Tajima’s *D* and heterozygosity. (L) Geographic distribution of pre-industrial pathogen stress. Circle colors indicate stress levels, with warmer colors representing higher pre-industrial pathogen stress and cooler colors lower stress. Arrow colors follow the color scheme used in Figure 3, and data points are color-coded by geographic regions. The solid line represents the best-fit linear regression, and dotted lines indicate non-significant associations. Shaded area shows the 95% confidence intervals.

A substantial portion of MHC-II variation was also explained by the temporal lag between the onset of agriculture and urbanization (Fig. 4D–E). Interestingly, this predictor, which reflects the speed of the transition between these major cultural phases and has not previously been considered in genomic studies, was significantly associated with both measures of genetic variation, explaining more than 22% of the variance (*R*^2^ = 0.274, *P* = 4.8 × 10^−5^ for Tajima’s *D* and *R*^2^ = 0.225, *P* = 2.8 × 10^−4^ for heterozygosity). In both independent regressions, shorter transition lags consistently corresponded to stronger signals of balancing selection in the MHC-II. The populations exhibiting these shorter transitions were broadly distributed and not restricted to a single geographic region (Fig. 4F).

The domestication-exposure *α* index was also significantly associated with the genetic measures in the independent regression analysis (Fig. 4G–H), explaining about 20% of their variance (*R*^2^ = 0.199, *P* = 7.1 × 10^−4^ for Tajima’s D; and *R*^2^ = 0.198, *P* = 7.3 × 10^−4^ for heterozygosity). The direction of the regressions was the same as in the multivariate models, with higher exposure to domesticated animals associated with higher signals of balancing selection. The populations with higher exposure and stronger signal of balancing selection are mainly from Europe, Middle East and Central Asia, while those with lower exposure to domesticated animals and weaker signals of balancing selection are mainly from America, Africa and Oceania (Fig. 4I).

The association between pre-industrial pathogen stress and genetic variation measures, which was significant for Tajima’s *D* but not for heterozygosity in the multivariate model for MHC-II (Fig. 3D), was not recovered in univariate analyses (Fig. 4J–L). This suggests that our pre-industrial pathogen stress variable does not explain MHC-II variation as a marginal predictor, but that its association with Tajima’s *D* becomes detectable after accounting for climatic conditions, long-term exposure to domesticated animals, and the timing of major cultural transitions.

To assess the robustness of these univariate regressions, we additionally performed an exhaustive sensitivity analysis by re-estimating the three significant associations after removing from the analysis all possible combinations of two and three populations (1,431 and 24,804 models, respectively). Across associations, 99% of the resampled models retained significance, supporting the robustness of the observed relationships (Fig. S8). Of the 26,235 resampled models, significance was lost in only four models, three of which involved the removal of the three Oceanian populations (Bougainville, the Papuan Highlands, and the Papuan Sepik). This suggests that these Oceanian populations strengthen the observed global patterns (while not being solely responsible for them).

Consistent with previous results, most of the independent relationships examined do not appear to be driven by the geographic distribution of populations in any consistent manner (Fig. 4), suggesting that the observed patterns are not simply the result of shared demographic history, but instead reflect specific selective pressures acting on immune-related regions.

## Discussion

We studied potential historical drivers of disease burden by developing an extensive dataset of historical-cultural, ecological, and climatic predictors, which we used to evaluate their association and relative contribution to global patterns of genetic variation in the MHC. The DisECCO dataset combines diverse data sources and extensive literature reviews to capture conditions that may have shaped disease burden during relevant historical periods. This approach is essential because current-day factors may only weakly reflect the historical conditions under which pathogen-mediated selection occurred. Using this dataset, we found that proxies for historical pathogen exposure, the lag between the onset of the Neolithic and the rise of urbanization, and climatic conditions were the strongest predictors of MHC genetic variation, particularly in MHC-II.

Among the environmental factors in the DisECCO dataset, the predictors that capture pathogen exposure are those most consistently associated with immune-related genomic signatures in previous studies [2, 22–24, 46]. Although high-resolution modern pathogen exposure data are available, current-day patterns are substantially affected by today’s socioeconomic, political, and health care system states and may only be weakly related to pathogen exposure levels during the evolutionary history of populations. Therefore, we compiled two predictors designed to reflect historical pathogen exposure levels, given available data. Our pre-industrial cumulative pathogen stress predictor was found to be positively associated with stronger signals of balancing selection and increased genetic diversity in different immune-related regions (Fig. 3A–B). Although the data we used were only about 100 years ago, many regions in the world have not yet transitioned to modern healthcare systems at that point in time. Therefore, at least for some geographic regions, these pathogen data may better reflect the disease burdens that preceded modern medicine interventions. However, we did not find a significant univariate association between pre-industrial pathogen stress and MHC-II genetic variation measures (Fig. 4J–L). We suggest that this absence of association reflects a conditional effect that emerges only after accounting for the influence of longer-term historical processes, such as colder Neolithic climatic conditions, shorter Δ*N* –*U*, or greater historical exposure to domestic animals, predictors that influence the burden of disease and are strongly associated with genomic signatures in our results.

Long-term contact with domesticated animals is a key pathway by which humans encountered novel pathogens. The spread of animal domestication has been closely tied to major transformations in human ecology and infectious disease dynamics, mainly through the zoonotic spillover of pathogens into human populations [24, 26, 47, 48]. Our index *α* aggregates the time populations have been in contact with each domesticated animal, as well the variety of domesticated animals documented in each population. We found that this predictor is positively associated with genetic variation in the MHC, particularly MHC-II (Fig. 3D), in both our multivariate and univariate analyses. Populations from Europe, the Middle East, and Central Asia showed both the highest levels of long-term exposure to domesticated animals and some of the strongest signatures of balancing selection in immune-related regions (Fig. 4I). These long-standing agricultural societies maintained close and continuous contact with a diverse array of domesticated animals over extended periods, likely intensifying pathogen exposure and shaping the genetic diversity of immune-related regions across populations. The east-west orientation of Eurasia may also have facilitated the diffusion of domesticated animals across similar latitudes and climatic conditions, increasing human exposure to zoonotic pathogens[49, 50].

Our analysis identified a previously underexplored predictor of immune-related genomic variation: the lag between the onset of the Neolithic and the rise of urbanization, which showed particularly strong associations with MHC-II variation (Fig. 3D and Fig. 4D–F). This result implies that shorter transition periods between these two major sociocultural shifts are associated with higher historical disease burdens and stronger balancing selection. Notably, this lag time explained much more of the variation in balancing selection signatures than the actual timing of the two transitions. One possible hypothesis explaining this result is that short lag times reflect a rapid sequence of transitions involving multifaceted change in multiple axes of living conditions, exposure to pathogens, population connectivity and disease transmission rates, which together synergized to generate a substantial and sustained increase in disease burden. Consistent with this interpretation, such peaks in disease burden have recently been observed in two main cultural centers in Eurasia by analyzing the increase in heterozygosity in ancient genomes [27]. In our dataset, several populations characterized by short Δ*N* –*U* values are associated with regions historically connected to major early centers of urbanization and exchange networks (Fig. 4F), including the Indus Valley sphere (Makrani, Balochi, Brahui, Sindhi, Pathan, and Hazara) and Eurasian steppe corridors (Uygur, Xibo, Daur, Yakut, and Russian). Although our analyses do not directly test the mechanisms underlying these regional patterns, they are consistent with the hypothesis that rapid transitions from agricultural to urban societies intensified pathogen exposure and contributed to stronger balancing selection in immune-related genomic regions.

A third factor that we identified as associated with genetic variation and balancing selection in the MHC, and most notably in MHC-II, is the climatic conditions at different points in time with respect to the cultural transitions in each population. We observe that colder and temperate climates, present-day and during Neolithic transitions and mainly in Central and Northeast Asia, were associated with stronger signals of balancing selection and higher levels of diversity in MHC-II. One possible explanation of this signal is that temperate climates, characterized by seasonal variation in pathogen exposure, may maintain multiple alleles over time, thereby favoring balancing selection, as different alleles can confer advantages during different seasons or years. In contrast, warmer and less climatically variable regions in the equatorial America, Africa, and Oceania were associated with lower levels of balancing selection, potentially reflecting a greater contribution of positive or purifying selection acting on immune-related genes. In these warmer and less variable environments, consistently high pathogen pressure can favor rapid fixation of advantageous resistance alleles, ultimately leading to reduced immune-related genetic diversity [51–53]. Another explanation for the higher disease burden signal in colder climates may be related to the living conditions in the cold season, which were characterized by living in enclosed and fairly crowded dwellings, often with domesticated animals. These conditions, which are conducive to high disease transmission and zoonotic spillover rates, may have been key drivers of increasing disease burden.

The DisECCO dataset combines information from diverse historical sources to approximate relevant conditions as closely as possible. However, the ability to link past ecological and cultural contexts to modern populations is limited and varies between populations. This limitation is expected to be affected by demographic processes, such as gene flow and population replacement, which can decrease the similarity between the modern population and ancient populations from the same region. Therefore, caution should be exercised when interpreting variables of specific HGDP populations. The uncertainties of our estimates require that robustness be ascertained while introducing substantial estimation errors (see Figs. S6–S7). However, noise in our variables would be expected to mask correlations with genomic signals rather than inflate them. Despite this noise, our results identified a number of significant correlations, suggesting that, on average, the variables meaningfully describe the ecological and cultural historical contexts of the HGDP populations.

Historical drivers of disease burden are difficult to reconstruct with precision. As a result, our predictors serve as proxies for the actual drivers of disease burden, and the interpretation of associations must be done with this in mind. Given the difficulty in characterizing cultural-ecological variables in past populations, our approach was to integrate factors such as domestication and cultural complexity over extended periods of time, further limiting the interpretation to broad patterns and associations; however, this also implies that positive and significant findings are dominant in driving disease burden levels. Additionally, several predictors are interrelated in complex ways. In particular, climatic variables show strong correlations ((|*r*| ≥ 0.8); Fig. S5), with models incorporating Neolithic climatic predictors often competing with similarly low AICc values against those including current climatic conditions or urbanization-period climatic conditions. However, including multiple climatic time points within the same model would introduce substantial collinearity. Although our statistical framework helps clarify some of these relationships, it cannot fully capture the multifaceted processes that shape the burden of the disease.

By pairing genomic data with historical-cultural, ecological, and climatic context, we show that immune-related genetic variation can be interpreted not only as a product of pathogen-mediated selection, but also as a record of the conditions that shaped pathogen exposure across human history. The strongest signals, particularly in MHC-II, point to the importance of major cultural transitions, long-term exposure to pathogens and domesticated animals, and climatic variation in shaping immune-related diversity among human populations. These results demonstrate the value of integrating genomic data with reconstructed historical context to understand how human cultural and environmental history has shaped the evolution of immune-related genomic variation.

## Methods

### Extracting immune-related genomic signatures

We extracted genomic signatures from the Major Histocompatibility Complex (MHC) and its three sub-regions. The MHC was defined as spanning positions 28,510,120 to 33,480,577 on chromosome 6, based on the Genome Reference Consortium Assembly Grch38.p14 (NCBI Map Viewer). Following the genomic coordinates proposed by Kulski *et al*. [54], we partitioned the MHC into three sub-regions: Class I from 29,723,434 (HLA-F) to 31,511,124 (MICB); Class III from 31,518,977 (PPIAP9) to 32,407,181 (BTNL2) and class II from 32,439,887 (HLA-DRA) to 33,131,343 (HLA-DPA3).

To quantify genetic diversity and identify signatures of selection in the MHC and its three sub-regions, we estimated heterozygosity (observed) and Tajima’s *D*, for the 54 populations of the HGDP dataset [34]. As a control, we performed the same calculations on the autosomal genome (chromosomes 1–22), excluding the MHC region. Heterozygosity was computed using VCFtools [55], first calculating per-individual values and then averaging them within each population. To control for genome-wide demographic effects and isolate MHC-specific heterozygosity, we first calculated heterozygosity for each population and then subtracted the corresponding genome-wide values from those of the MHC and its three sub-regions (Δ*H_o_*).

Tajima’s *D* was also estimated using VCFtools, with a sliding window approach (window length: 1000 bp; step size: 1000 bp) applied per population (non-overlapping). To identify significant deviations from neutrality, we retained only windows with Tajima’s *D* values greater than +2 or less than −2 [56], and excluded those with fewer than 10 segregating sites to minimize false positives. Mean Tajima’s *D* values were calculated per population using the remaining significant windows.

To compare heterozygosity and Tajima’s *D* between the MHC and its sub-regions, we used paired Wilcoxon signed-rank tests across the 54 HGDP populations.

### Assessing the relationship between immune-related genomic signatures and distance from Africa

To evaluate whether genetic variation in the MHC and its three sub-regions among the 54 HGDP populations is shaped by the same demographic processes underlying genome-wide variation, we computed Pearson’s correlation coefficients (*r*) and performed linear regression analyses in R using the cor() and lm() functions, respectively. We first compared immune-related and genome-wide signatures with the geographic distance from Africa of each population, used as a proxy for neutral demographic patterns associated with human migration history [20, 57, 58]. Then we assessed the relationship between immune-related and genome-wide signatures directly to determine the extent to which MHC patterns reflect genome-wide demographic variation.

Geographic distance from Africa was estimated for all populations outside southern and central Africa following the methodology of Ramachandran *et al.* [20]. Distances were calculated using the haversine formula, assuming an Earth radius of 6,371 km and incorporating six waypoints to approximate human migration routes. These waypoints included Cairo, Egypt (30N, 31E); Istanbul, Turkey (41N, 29E); Phnom Penh, Cambodia (11.5N, 105E); Anadyr, Russia (64.5N, 177.5E); Nome, Alaska, USA (64.5N, 165.4W); and Mexico City, Mexico (19.5N, 99W). Calculations were implemented using a custom R function (scripts available at https://github.com/Greenbaum-Lab/MHC_Analysis). Populations from southern and central Africa, including Mandenka, Yoruba, Biaka, Mbuti, Bantu Kenya, San, and Bantu South Africa, were treated as origin centers and assigned a distance of zero. The corresponding values for this predictor are available in *Supplementary Dataset S6*.

### Compiling the DisECCO dataset

To understand the inter-population variation in the immune-related regions, we developed DisECCO, a dataset of historical–cultural, ecological, and climatic predictors potentially associated with the burden of infectious diseases throughout history (Fig. 2).

#### Onset of Neolithic revolution and Urbanization

The onset of the Neolithic Revolution and the rise of urbanized societies, characterized by unprecedented spatial and social agglomeration, represent two major sociocultural transitions that have significantly shaped the burden of infectious diseases throughout human history [59]. These transitions occurred at different times across geographic regions and populations. We extracted information on the timing of these two transitions from the Seshat databank [40], specifically the “Agricultural Productivity” sub-dataset [60], and from the ArchaeoGLOBE database [41]. Because these databases are organized by regions rather than HGDP populations, each population was assigned the values corresponding to the closest matching region. Since no single source provides complete coverage for all populations, these databases were complemented with information from the literature when available (i.e., [61–66]). Detailed information on population matching and the relevant literature for this predictor is provided in *Supplementary Dataset S1*.

#### Time lag between the onset of the Neolithic and urbanization

The transition from the Neolithic Revolution to the emergence of urbanization may have represented a continuous process of increasing social complexity or, in some regions, two distinct historical transitions separated by a long period of time. The temporal interval between these two major transitions may therefore reflect different trajectories of sociocomplexity growth and may have influenced disease burden in different ways. To evaluate this possibility, we included the absolute difference between the estimated onset of Neolithic and urbanization, Δ*N* –*U* for each population as a predictor to our dataset.

#### Sociopolitical complexity

During the past 10,000 years, human societies have undergone significant cultural transformations. The evolution of sociocultural variables such as hierarchical structures, territorial organization, infrastructure, and information systems has potentially facilitated increases in population density and, consequently, the establishment and transmission of pathogens. Turchin *et al.* [42] quantified social complexity by collecting 51 variables describing cultural and sociopolitical development in 30 natural geographic areas over the past 10,000 years and suggested that PC1 of this dataset captures much of the variation in social complexity across these regions. We summarized the PC1 trajectory over time by calculating the area under the curve (AUC) for each of the 30 regions using the ‘Trapz’ function of the R package caTools [67]. Script used for this analysis are available at https://github.com/Greenbaum-Lab/MHC_Analysis. The AUC of the geographically closest region was then assigned to each of the 54 HGDP populations as a measure of long-term sociopolitical complexity. The values for each population and information on population matching are provided in *Supplementary Dataset S2*.

#### Current and ancient climate

Disease ecology, pathogenic environments, and human behavior and physiology are all affected by climatic conditions [38]. Therefore, considering climatic conditions, particularly those associated with major episodes of changes in pathogen exposure, is important in order to explain variation in disease burden. We gathered climatic data corresponding to the geographic location associated with each population for the present day, as well as for the periods marking the onset of the Neolithic and urbanization. Current climatic variables were obtained from the CHELSA V2.1 bioclimatic dataset [68]. Past climatic conditions were extracted from the CHELSA-traCE21k database [43]. Both datasets provide 19 bioclimatic variables, with CHELSA-traCE21k covering climatic conditions across the Earth’s surface over the last 20,000 years. We sampled the rasters at 1,000-year intervals, starting from the present and extending back to 20,000 years before present. To reduce computational demands, 1 km cells were aggregated to a resolution of ∼100 km using the R package *terra* [69], and the mean value of each bioclimatic variable was calculated for each aggregated cell. We then performed a principal component analysis (PCA) to summarize the climatic variation on Earth’s surface over time. The first two components captured the main axes of climatic variability, with PC1 explaining 52% and PC2 explaining 18% of the total variance. The four main contributors to PC1 were temperature-related variables, whereas three of the four main contributors to PC2 were precipitation-related variables (Fig. S4). From these PCs, we extracted climatic values for each HGDP population at three time points: (i) the time slice closest to the estimated onset of the Neolithic Revolution (see *Onset of Neolithic revolution and Urbanization* above), (ii) the time slice closest to the estimated onset of urbanization, and (iii) the present day. This approach captures key dimensions of climatic variation across the HGDP populations. The corresponding values for these predictors, along with the geographic coordinates used in this analysis, are provided in *Supplementary Dataset S3*. The script used to process the climatic data and derive these predictors are available at https://github.com/Greenbaum-Lab/MHC_Analysis.

#### Exposure to domesticated animals (α index)

Domestication of animals is another major sociocultural process that has influenced human societies throughout history, increasing the risk of zoonotic pathogen transmission due to enhanced contact between humans and animals [26, 47, 48]. The extent of this influence varies between populations depending on both the diversity of domesticated species and the historical timing of their domestication. To assess this effect, we compiled information from the literature on the timing of domestication events for 15 key species in the geographic regions associated with the 54 HGDP populations (see [70, 71] for reviews), including chickens, horses, donkeys, buffaloes, pigs, cows, goats, sheep, cats, dogs, llamas, alpacas, turkeys, guinea pigs, and mice. We then developed an index, *α*, for each population that quantifies total exposure to domesticated animals. This index is defined as:

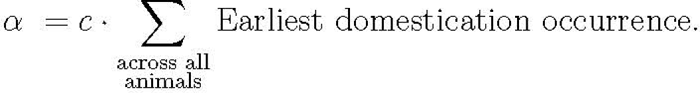

where *c* is a normalizing constant equal to one over the product of the number of animals (15) and the age of the oldest domestication event (14,500), i.e. 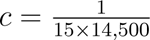. This index quantifies cumulative exposure to domesticated animals over time, incorporating both the diversity of species and the timing of their domestication. Further details on this predictor, the associated bibliography, and the estimated *α* values are provided in *Supplementary Dataset S4*.

#### Pre-industrial pathogen stress

To gather historical information on pre-industrial pathogen stress for each population, we used the D-PLACE database[44], specifically the standard cross-cultural sample dataset (SCCS) [72, 73] and the Total Pathogen Stress predictor [SCCS1260]. The SCCS dataset contains pre-industrial records on the prevalence levels of seven pathogens (Leishmania, trypanosome, malaria, schistosome, filariae, spirochetes, and leprosy) in 186 human populations [74]. Some of these data were collected more than 100 years ago, minimizing the influence of current differential access to modern healthcare on the prevalence of pathogens. We matched each of our 54 populations to the closest non-industrial population in D-PLACE to create this variable. Detailed information on population matching and the corresponding predictor values is available in *Supplementary Dataset S5*.

### Assessing the relationship between immune-related genomic signatures and DisECCO predictor variables

To explore the factors contributing to inter-population differences in MHC and its three sub-regions, we conducted multivariate regression analyses, as a control we also re-ran the analysis on genome-wide variation. Each historical–cultural, ecological, or climatic variable was standardized to z-scores (mean = 0, standard deviation = 1) using the ‘scale’ function in the R package *base* [75]. To select the best multiple model with the smallest number of predictor variables, we combined the LASSO method [76] with the Akaike Information Criterion (AICc [77]), following [78–80]. Specifically, we applied LASSO variable selection, using the ‘cv.glmnet’ function of the glmnet R package [81], with the penalty parameter *λ* chosen by cross-validation analysis. Given the high dimensionality of our data, we further refined the set of variables selected through LASSO by identifying the most parsimonious and interpretable best–fitting models using the AICc criterion with the ‘dredge’ function of the MuMIn R package [82]. The best model was defined as the one with the lowest AICc [45]. Finally, to evaluate the effect of multicollinearity among predictors in the best-fitting models, we calculated the mean variance inflation factor (*V IF*) across predictors using the ‘vif’ function from the car R package [83].

The DisECCO dataset for the HGDP populations is inherently prone to estimation errors. To assess the robustness of our best multiple models to potential inaccuracies, we performed a sensitivity analysis by progressively introducing random errors of 10%, 20%, 30%, and 40% into the 13 predictors. For each error level, 100 random perturbations were generated, and best–fitting models were re-identified as described above. A predictor variable was considered robust if it was retained in at least 50% of the best–fitting models under up to 20% random error (variable that were not included in the original best–fitting model without error were not considered even if they were included when error was introduced). Higher error levels (30% and 40%) were included to characterize the overall stability of predictors beyond the robustness threshold, allowing a visual comparison of how rapidly different variables lose explanatory power under increasing uncertainty. This sensitivity analysis was implemented using a custom R function developed specifically for this study (script available at https://github.com/Greenbaum-Lab/MHC_Analysis). As a baseline comparison, we repeated the procedure 100 times without introducing any errors. Minor variations in predictor retention in the error-free runs were attributed to stochasticity in the LASSO model fitting procedure, specifically due to differences in the cross-validation folds used to select the penalty parameter (*λ*). These replicates therefore provide a baseline level of variability against which the effect of introducing random errors can be evaluated.

To calculate the proportional contribution of the robust variables to the overall slopes in each best–fitting model, we first standardized the predictor variables and fit a linear model in which the response variable was regressed on the standardized predictors. We then extracted the regression coefficients and normalized each by dividing it by the sum of the absolute values of all coefficients. Finally, we calculated the percentage contribution of each variable to the model’s *R*^2^ by multiplying its normalized contribution by the *R*^2^ value.

To further evaluate the independent effects of the main predictors identified in the multivariate analyses, we conducted univariate regression analyses focusing on the four predictors associated with MHC-II variation: (i) climatic conditions during either the current (for Tajima’s *D*) or Neolithic (for heterozygosity) periods, (ii) pre-industrial pathogen stress, (iii) long-term contact with domestic animals (*α* index), and (iv) the temporal lag between the onset of agriculture and urbanization (Δ*N* –*U*). To evaluate the robustness of significant univariate associations to influential populations, we performed an exhaustive sensitivity analysis by re-estimating each significant regression after removing all possible combinations of two and three populations (1,431 and 24,804 models, respectively). For each resampled dataset, the regression was refitted and the significance of the association was recorded. This procedure allowed us to assess whether the observed relationships were consistently supported across population subsets or disproportionately influenced by a small number of populations.

## Data Accessibility

All datasets supporting this study are openly available on GitHub at https://github.com/Greenbaum-Lab/DisECCO-dataset. These include: (i) Dataset S1, Ages of the onset of the Neolithic revolution and urbanization for HGDP populations; (ii) Dataset S2, PCA-derived climatic variables across temporal slices for HGDP populations; (iii) Dataset S3, Area under the curve of sociopolitical complexity trajectories for HGDP populations; (iv) Dataset S4, Ages of the oldest domestication events for HGDP populations; (v) Dataset S5, Pre-industrial pathogen stress for HGDP populations; (vi) Dataset S6, full DisECCO dataset: Comprehensive set of historical, cultural, ecological, and climatic predictors compiled to explain variation in immune-related genomic signatures across the 54 HGDP populations. The R scripts used for these analyses are available at https://github.com/Greenbaum-Lab/MHC_Analysis.

## Acknowledgments

This work was supported by the Minerva Center for the Study of Population Fragmentation (MCPF).

## Author contributions

Conceptualization: AMG, KH, GG; Study design: AMG, KH, NM, GG; Coding: AMG; Analyses: AMG, GG; Writing of Original Draft: AMG; Review and Editing: AMG, KH, NM, GG; Funding Acquisition: AMG, GG.

