## Supplementary Figures and Tables for "Drivers of immune-related genetic variation across human populations"

### 1 Supplementary Figures

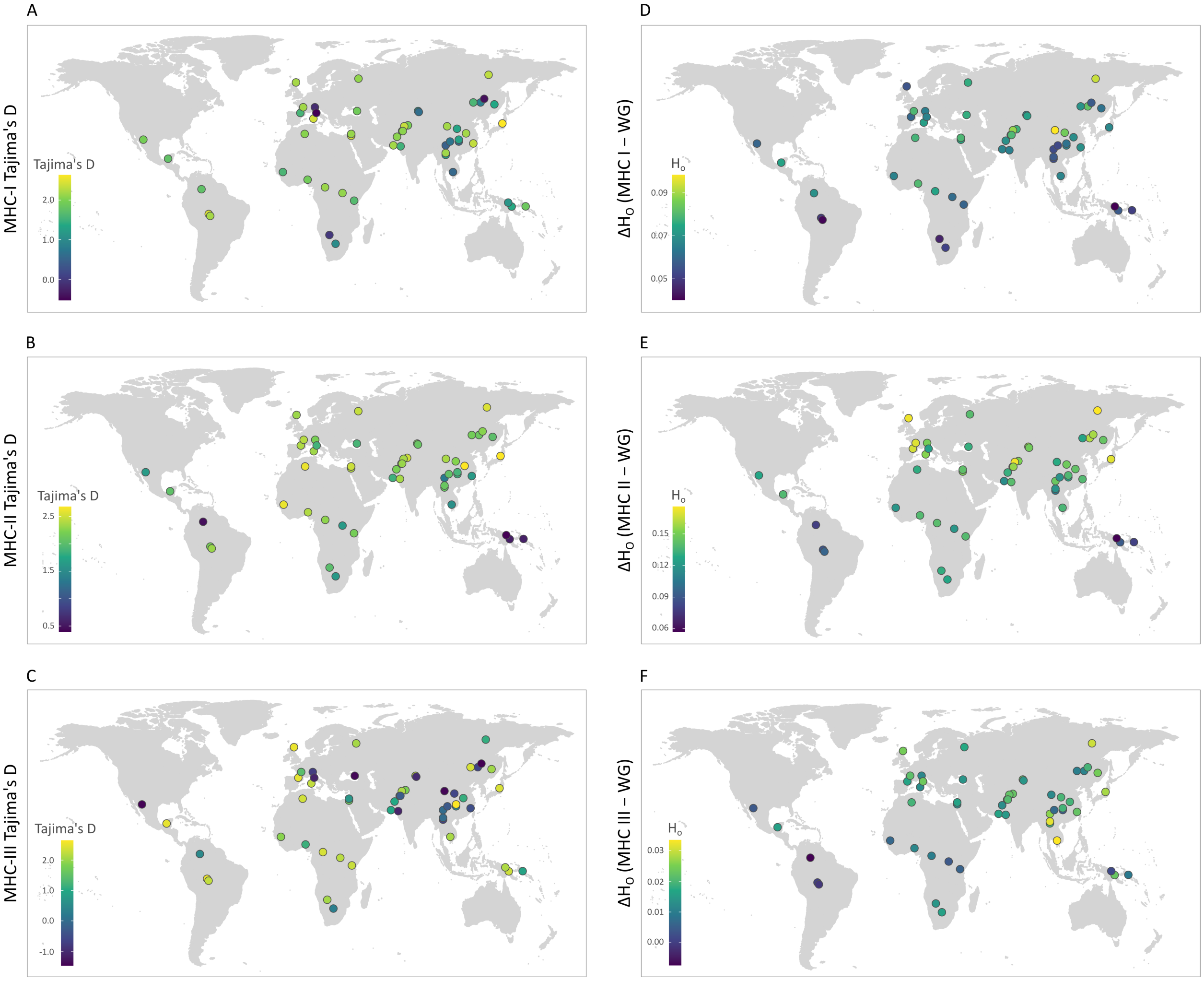


Figure S1: **Worldwide patterns of genomic variation in the MHC sub-regions across 54 human populations.** (A–C) Tajima's $D$ for the MHC-I, MHC-II, and MHC-III, respectively. (D–F) Heterozygosity for the MHC-I, MHC-II, and MHC-III, respectively.

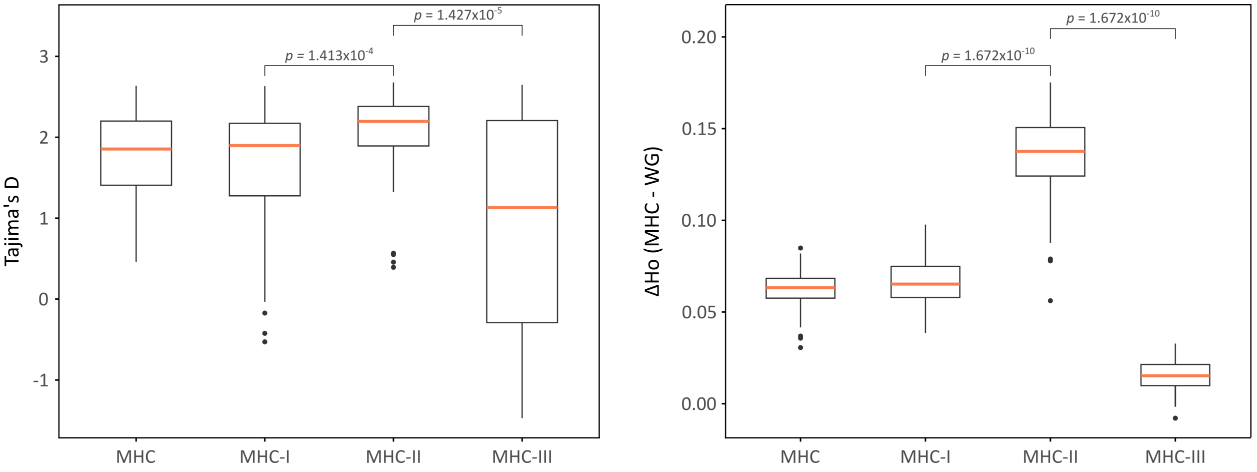


Figure S2: **Comparison of genetic variation among MHC classes across 54 HGDP populations.** (A) Distribution of Tajima's $D$ values for the complete MHC region and its three major functional classes (MHC-I, MHC-II, and MHC-III). (B) Distribution of heterozygosity for the same regions. Boxplots show the median, interquartile range, and values within 1.5 times the interquartile range; points represent outliers. *P-values* correspond to paired Wilcoxon signed-rank tests comparing MHC-II with MHC-I and MHC-III. Across populations, MHC-II exhibited significantly higher Tajima's $D$ and heterozygosity values than both MHC-I and MHC-III, consistent with stronger signatures of balancing selection and elevated genetic diversity in this region.

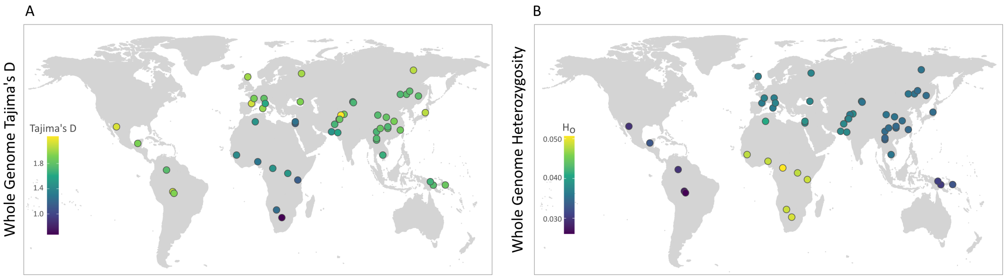


Figure S3: **Worldwide genome-wide patterns of variation across 54 human populations.** (A) Genome-wide Tajima's $D$. (B) Genome-wide heterozygosity.

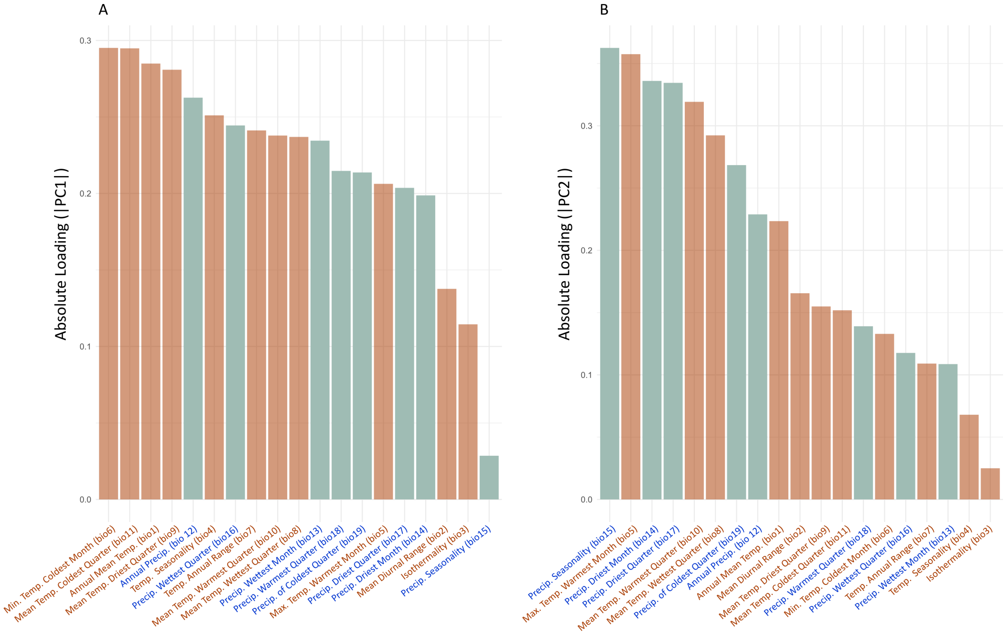


Figure S4: **Contribution of bioclimatic predictors to the first two principal components from a PCA of 19 bioclimatic variables.** The y-axis shows the absolute value of the correlation coefficient (loading) between each variable and the corresponding principal component: (A) PC1 and (B) PC2. Red bars represent temperature-related variables, and blue bars represent precipitation-related variables.

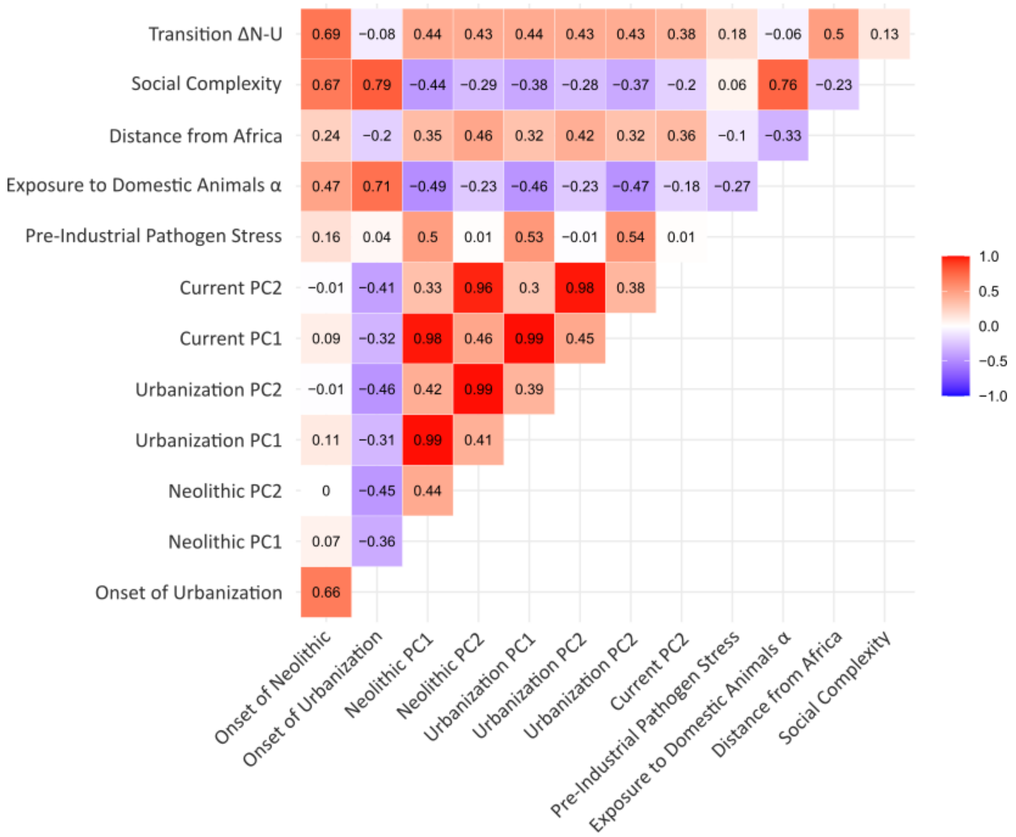


Figure S5: **Pairwise correlations among variables in the DisECCOdataset.** Shown are pairwise Pearson correlation coefficients among the 13 predictors used in the analysis: onset of the Neolithic and urbanization; PC1 and PC2 from climatic variables of the Neolithic, urbanization, and current climate; pre-industrial pathogen stress; $\alpha$ index; distance from Africa; sociopolitical complexity; and the time lag between Neolithic onset and urbanization. Colors indicate the strength and direction of Pearson correlation coefficients, ranging from $-1$ (strong negative correlation) to 1 (strong positive correlation). Correlation coefficients are displayed within each cell. Correlations with an absolute value greater than 0.7 were considered strong.

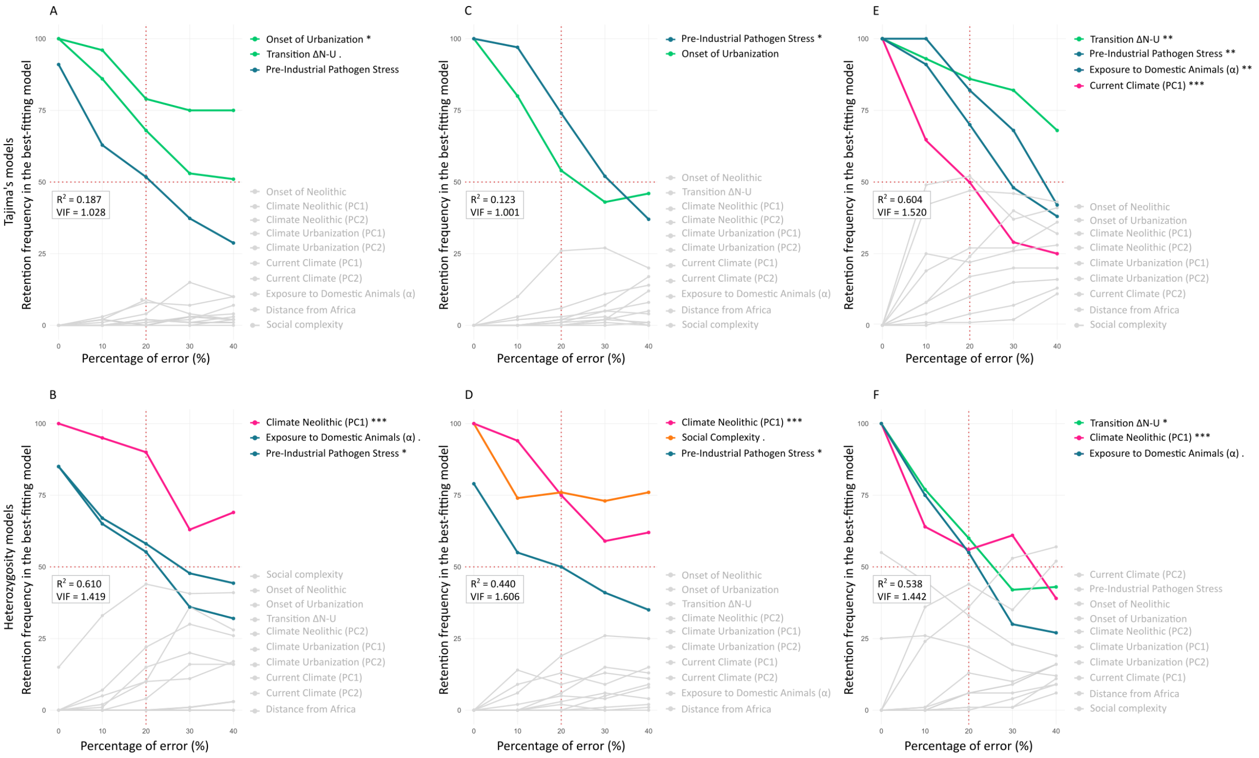


Figure S6: **Sensitivity analysis of predictor robustness for the best-fitting models of MHC, MHC-I, and MHC-II genomic signatures after introducing random errors of 10%, 20%, 30%, and 40% into the 13 predictor variables.** (A) Frequency with which each variable is retained in the best–fitting model of MHC Tajima's $D$, relative to the percentage of random errors introduced. Dark colors highlight variables that passed the robustness criterion, defined as appearing in at least 50% of the best–fitting models under up to 20% random error, provided that the variable was included in the original best–fitting model without error. (B) Frequency of variable retention in the best–fitting model of MHC heterozygosity. (C–D) Frequency of variable retention in the best–fitting models of MHC-I Tajima's $D$ and heterozygosity. (E–F) Frequency of variable retention in the best–fitting models of MHC-II Tajima's $D$ and heterozygosity. $R^2$ and $VIF$ and coefficient significance levels shown in each panel correspond to the best-fitting model composed only of predictors that met the robustness criterion. Asterisks indicate the significance level of the estimated coefficient for each predictor: '.' $p < 0.1$ (marginally significant), '*' $p < 0.05$ (significant), '**' $p < 0.01$ (very significant), and '***' $p < 0.001$ (highly significant).

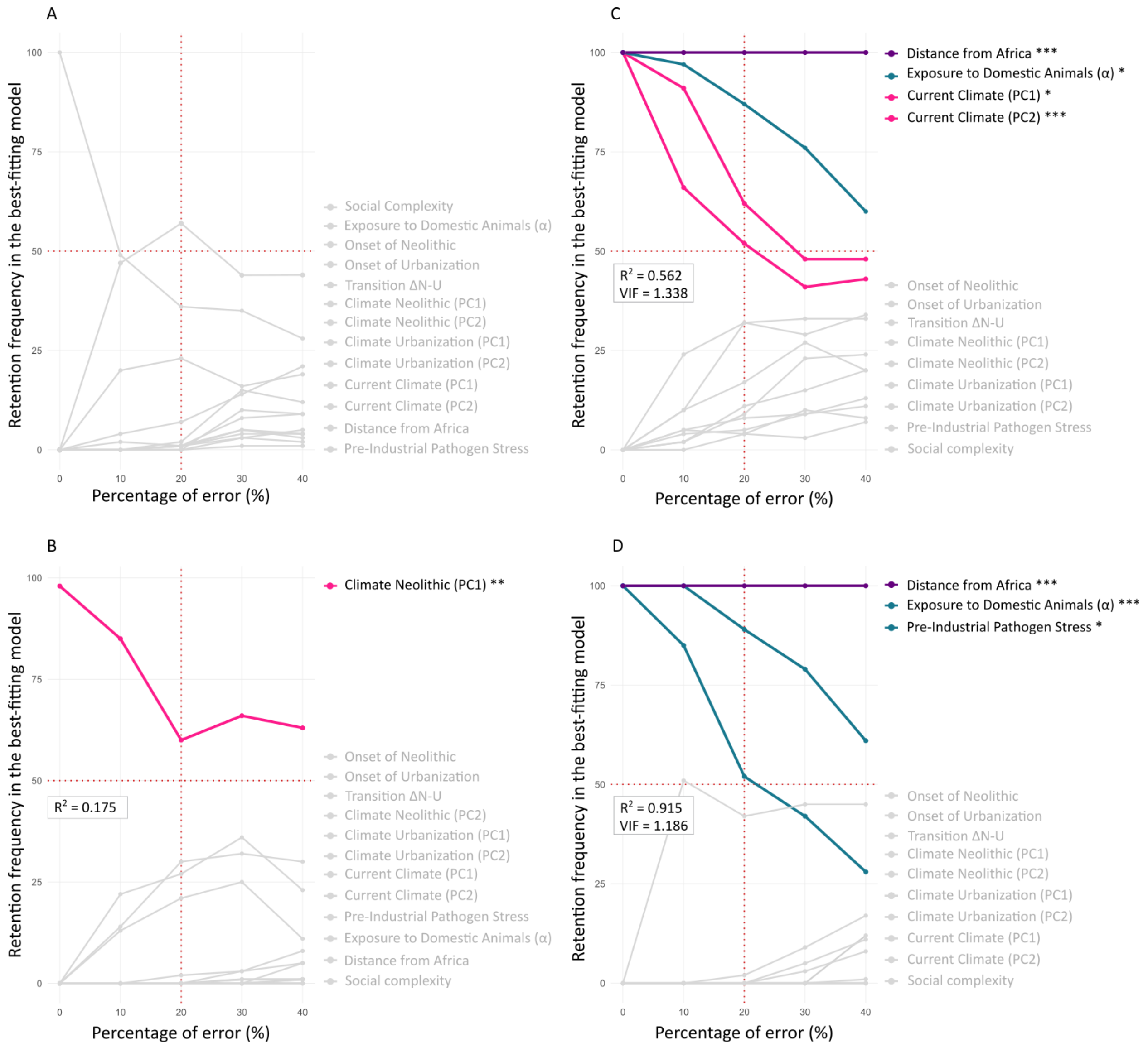


Figure S7: **Sensitivity analysis of predictor robustness for the best-fitting models of MHC-III and genome-wide genomic signatures after introducing random errors of 10%, 20%, 30%, and 40% into the 13 predictor variables.** (A) Frequency with which each variable is retained in the best–fitting model of MHC-III Tajima's $D$, relative to the percentage of random errors introduced. (B) Frequency of variable retention in the best–fitting model of MHC-III heterozygosity. Dark colors highlight variables that passed the robustness criterion, defined as appearing in at least 50% of the best–fitting models under up to 20% random error, provided that the variable was included in the original best–fitting model without error. (C–D) Frequency of variable retention in the best–fitting models of genome-wide Tajima's $D$ and heterozygosity. $R^2$ and $VIF$ and coefficient significance levels shown in each panel correspond to the best-fitting model composed only of predictors that met the robustness criterion. Asterisks indicate the significance level of the estimated coefficient for each predictor: '.' $p < 0.1$ (marginally significant), '*' $p < 0.05$ (significant), '**' $p < 0.01$ (very significant), and '***' $p < 0.001$ (highly significant).

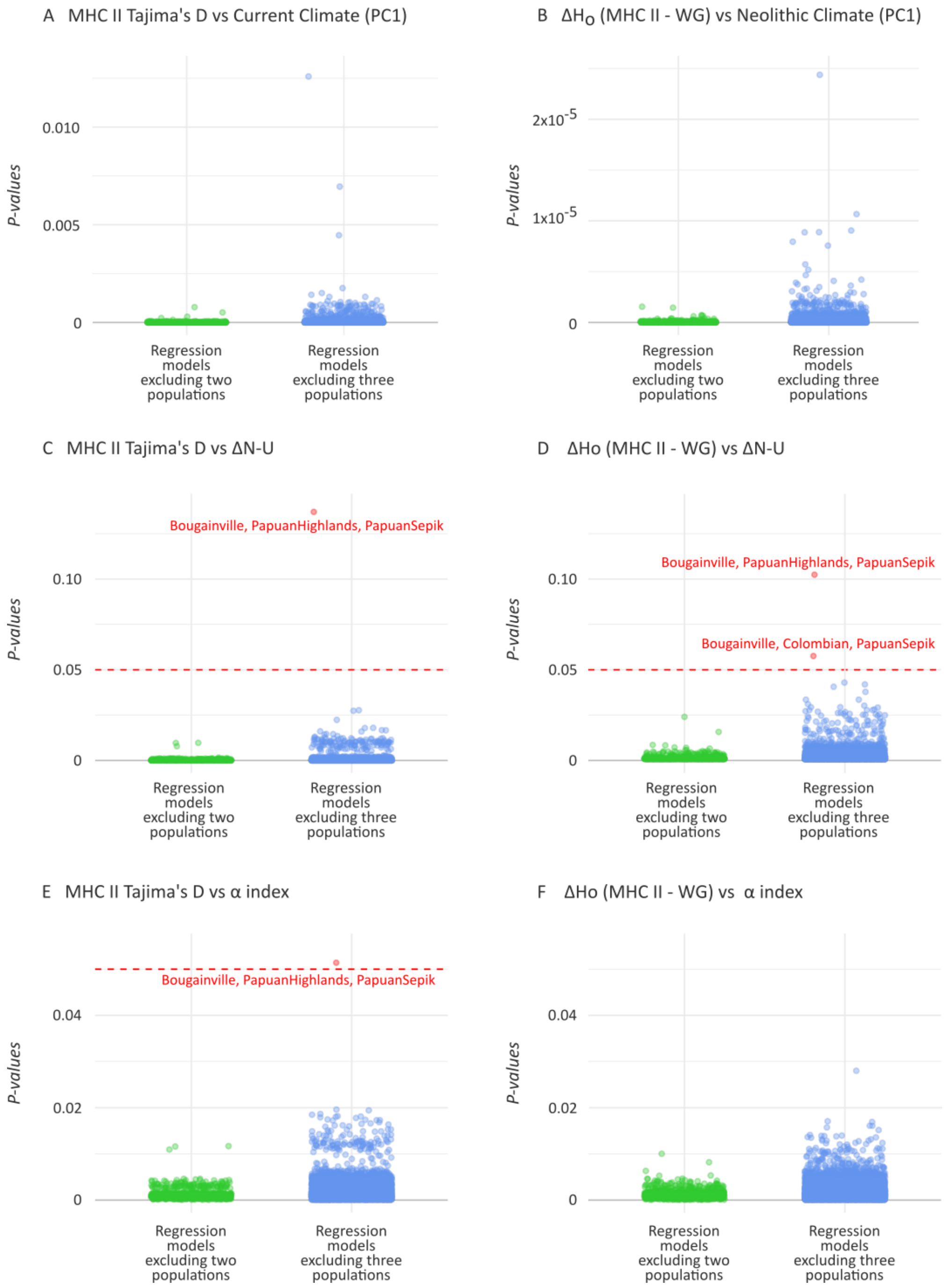


Figure S8: **Sensitivity analysis evaluating the robustness of the three significant univariate associations identified for MHC class II.** For each association, the regression was re-estimated after removing all possible combinations of two populations (green; 1,431 models) and three populations (blue; 24,804 models). Panels show the distribution of resulting p-values for the associations between MHC-II Tajima's $D$ and current climate (PC1) (A), heterozygosity and current climate (PC1) (B), MHC-II Tajima's $D$ and the lag between the onset of the Neolithic and urbanization ($\Delta N$–$U$) (C), heterozygosity and $\Delta N$–$U$ (D), MHC-II Tajima's $D$ and long-term exposure to domesticated animals ($\alpha$) (E), and heterozygosity and $\alpha$ (F). The red dashed line indicates the significance threshold ($p = 0.05$). Across all resampled models, significance was retained in more than 99% of cases.

### 2 Supplementary Tables

Table S1: Linear regression analyses between genomic signatures (Tajima's $D$ and heterozygosity) and distance from Africa

| Signatures | Tajima's D | | Heterozygosity | |
|---|---|---|---|---|
| | $r$ | *P-value* | $r$ | *P-value* |
| GW | 0.558 | $1.19 \times 10^{-5}$ | -0.882 | $1.20 \times 10^{-18}$ |
| MHC | -0.144 | 0.299 | -0.389 | $3.63 \times 10^{-3}$ |
| MHC-I | 0.060 | 0.666 | -0.261 | $5.64 \times 10^{-2}$ |
| MHC-II | -0.417 | $1.72 \times 10^{-3}$ | -0.421 | $1.51 \times 10^{-3}$ |
| MHC-III | 0.005 | 0.971 | -0.212 | $1.23 \times 10^{-1}$ |

Numbers shown are the Pearson correlation coefficients ($r$) and *P-values* for the association between Distance from Africa and Tajima's $D$/heterozygosity across different genomic regions. $R^2$ values (variance explained) can be obtained by squaring $r$. For example, $r = -0.882 \rightarrow R^2 \approx 0.777$.GW = Genome-wide signatures.

Table S2: Correlations between Tajima's $D$ Signatures

| Signatures | GW | MHC | MHC-I | MHC-II | MHC-III |
|---|---|---|---|---|---|
| GW | 1.000 | | | | |
| MHC | n.s. | 1.000 | | | |
| MHC-I | n.s. | 0.856 | 1.000 | | |
| MHC-II | n.s. | 0.670 | 0.274 | 1.000 | |
| MHC-III | n.s. | 0.427 | 0.339 | n.s. | 1.000 |

Numbers shown are the Pearson correlation coefficients ($r$). Only significant results ($p \leq 0.05$) are shown. "n.s." = not significant. GW = Genome-wide signatures.

Table S3: Correlations between Heterozygosity Signatures

| Signatures | GW | MHC | MHC-I | MHC-II | MHC-III |
|---|---|---|---|---|---|
| GW | 1.000 | | | | |
| MHC | n.s. | 1.000 | | | |
| MHC-I | n.s. | 0.804 | 1.000 | | |
| MHC-II | n.s. | 0.894 | 0.506 | 1.000 | |
| MHC-III | n.s. | 0.601 | 0.245 | 0.574 | 1.000 |

Numbers shown are the Pearson correlation coefficients ($r$). Only significant results ($p \leq 0.05$) are shown. "n.s." = not significant. GW = Genome-wide signatures.
